# Mortality and malformation effects of boric acid against larval stage of *Aedes aegypti* (Culicidae: Diptera)

**DOI:** 10.1101/2023.09.05.556246

**Authors:** Somia Eissa Sharawi

## Abstract

Mosquitoes, the disquieting insect, can transmit pathogens more than any other insect, and they can cause fatal diseases to both humans and animals in their environment. With the increasing problem of pesticide resistance spreading in the field, substantial efforts are being made at the international level to develop environmental insecticides to reduce these mosquito populations or limit their potential to transmit disease. In this preliminary study, mortality and malformation effects of seven concentrations (700, 1000, 1200, 1500, 2000, 2500, and 3000 ppm) of boric acid were evaluated against the 4^th^ larval stage of *Aedes aegypti* after 24 h of exposure. Malformed and dead larvae were photographed under the microscope with a digital camera connected to a computer stereo-dissecting microscope. Our results showed that the lethal concentrations (LC_50_ and LC_90_) were 1312.49 and 2578.26, respectively. Also, the results showed malformation effects against treated larvae with boric acid compared to the control group. These malformations manifested in albino pupa, Pigmentation, Larval-pupal intermediate, Black coloration at the posterior end, Opaque swelling on the thorax, Segment contraction, Deformed cuticles, black coloration at the posterior end, and nick elongation. Boric acid is less toxic than synthesized insecticides, and in low concentrations, it causes mortality and malformation effects against larval stages of *A. aegypti*. Further, the use of boric acid as a safe method for controlling larval stages of *A. aegypti* can be recommended as it is available, cost-effective, and less harmful to humans, animals, and the environment.

## Introduction

Mosquitoes are considered one of the deadliest insects for humans and animals due to the dependence of their females on the blood to obtain their food. This parasitism behavior has supported the ability of these insects to transmit many serious and widespread pathogens between vertebrate hosts, including the most important human pathogens: Malaria, Dengue fever, and yellow fever. *Aedes* species were the most mosquitoes found in Jeddah, Saudi Arabia ^[1]^. According to new research from the Middle East focusing on Dengue disease, there is a remarkable increase in this disease, particularly in the Kingdom of Saudi Arabia, and controlling methods were used against these annoying insects using conventional insecticides ^[2]^. Chemical control was and still is the most popular and effective way, but its control as insecticides is not suitable for several reasons ^[3]^. This chemical shows several environmental and health effects on plants, animals, and humans. Unfortunately, mosquitoes are developing resistant insects to conventional insecticides ^[4]^. For this reason, many research studies focused on different resources and applications as a safe way to control them without affecting non-target organisms. Inorganic compounds such as boric acid are natural materials. Usually, boric acid is mined from the mineral sassolite, and it can control pests ^[5]^. Boric acid has a long history as an insecticide in pest management, and it is an effective alternative to conventional neurotoxic insecticides. ^[6]^ confirmed that boric acid was a safe alternative insecticide because of its inorganic compound with low toxicity and non-volatile mineral. The boric acid mode of action against insects is not clear yet, although the destruction of the digestive tract wall and penetration of the exoskeleton has been reported; also, the subject of many researchers and several modes of action have been proposed, such as abrasion of the cuticle, the neurotoxic action or disruption of the midgut ^[7]^. Many studies of boric acid were determined as an adult stage mosquito control agent, not the larval stage. ^[8]^ found that boric acid significantly decreases the Acetylcholinesterase activities in some insects. ^[9]^ found that cockroaches and other insects were controlled by boric acid. Much of the work regarding boric acid efficacy has been done on the adult stages of some insects including mosquitoes; however, very little data is available concerning mosquitos’ larval stage. Therefore, the present work was designed to identify the mortality and malformation effects of boric acid against mosquitos’ larval stage as a safety method for control and find the possibility of a new method to control.

## Materials and Method

### Tested mosquitoes

The experiments were carried out at Dengue Mosquito Research Station in King Abdulaziz University, Jeddah (Saudi Arabia). Lab strains of the 4^th^ larval stages were used in the study and kept under laboratory conditions (27± 3 °C, 70±5 % relative humidity (R.H.), and 12:12 light to dark (L:D)). Larvae stages were fed on fish food in equal amounts (1:1). Newly formed pupae were transferred to a new plastic cup containing water for further experiments.

### Boric acid preparation

Boric acid was purchased from AL-Shafei Medica & Scientific Equipment Exh. Jeddah, Saudi Arabia, and used during bioassay. The choice of these inorganic compounds was based on the fact that these insecticides are available for everyone, low coast, and have not much tested against larval stages of *A. aegypti* in Jeddah province. A stock solution of boric acid was prepared by adding 1 ml of boric acid to 99 ml of distilled water and ensuring that boric acid was completely soluble in water. Seven concentrations of boric acid (700, 1000, 1200, 1500, 2000 2500, and 3000 ppm) were prepared according to the following formula and used for further experiments.

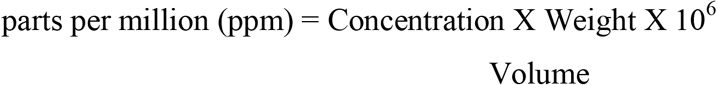

### Larval bioassay

The susceptibility of *A. aegypti* larvae was handled according to the method of ^[10]^ Treatments were continued for 24 hours by exposing the 4^th^ instar to the concentrations of boric acid. Five replicates of 20 larvae per concentration were used in 200 ml plastic cups containing 100 ml tap water. Tab water was only used in the control group. Abnormalities and mortalities of larvae and pupae were recorded every 4 hours. The biological effects of boric acid were recorded as the percentage of the larval stage that does not move and develop into a pupa. Malformed and dead larvae were photographed under the microscope with a digital camera connected to a computer stereo-dissecting microscope.

### Data Analysis

This study was conducted in a completely randomized design (CRD). The study was based on the 4^th^ larval stages of *A. aegypti*. LC_50_ and IC_50_ were calculated according to the Probit analysis program with the lower and upper confident limits and the inclination of the toxicity line and Chi-square ^[11].^ All the experiments photographs were taken by dissecting microscope with a digital camera. If the mortality control was between 5% and 20%, the mortalities of treated groups were corrected according to Abbott’s formula ^[12]:^

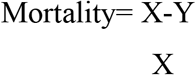

Where X = percentage surviving of untreated insects (control) and Y= percentage surviving of treated insects.

## Results and Discussion

Our study showed the mortality and malformation effects of boric acid against the 4^th^ of larval stage of *A. aegypti*, treated with a series of concentrations after 24 h of exposure. The general mortality of *A. aegypti* was significantly different from the control (Table 1), and boric acid was found to be effective in the suppression of *A. aegypti*. The effective concentrations were between 1500 −3000 ppm, and the corresponding mortality rate of these concentrations was between 60 – 94 %. Boric acid is a safe material for the control of the mosquito larvae, which was confirmed by the LC_50_ and LC_90_ values (1312.49 −2578.26), respectively (Table 1 and Fig. 1). This agrees with ^[13]^ who studied the mortality of boric acid against *A. aegypti*. Also, ^[14]^ reported that 1% boric acid amalgamated with sugar was an effective bait to kill adult mosquitoes. ^[15]^ investigated that 1% of boric acid caused 100% mortality against some *Aedes* species eggs and the effective concentration was 1% for the oviposition attraction (0.5%).

**Table 1:** Mortality effect of boric acid against *A. aegypti* larvae after 24 h.

| **Mortality** |  |  |  |
| --- | --- | --- | --- |
| **Con. (ppm)** | **After 24 h.** | **Linear response %** | **Linear probit** |
| 700 | 17 | 11.6473 | 3.8068 |
| 1000 | 28 | 30.2891 | 4.4838 |
| 1200 | 40 | 43.2515 | 4.8300 |
| 1500 | 56 | 60.0080 | 5.2535 |
| 2000 | 77 | 78.7950 | 5.7994 |
| 2500 | 89 | 88.9302 | 6.2229 |
| 3000 | 99 | 94.1582 | 6.5691 |
| Control | 0 | 0 | 0 |
| LC<sub>50</sub><br>(L. limit – U. limit) | 1312.49<br>(1237.390-1387.42) |  |  |
| LC<sub>90</sub><br>(L. limit – U. limit) | 2578.26<br>(2372.69- 2854.79) |  |  |
| Slope | 4.3706 |  |  |
| Tabulated (Chi)<sup>2</sup> | 11.1 |  |  |
| Calculated (Chi)<sup>2</sup> | 6.6975 |  |  |

When tabulated (Chi)<sup>2</sup> larger than calculated at 0.05 level of significance indicates the
homogeneity of results

**Fig 1:**
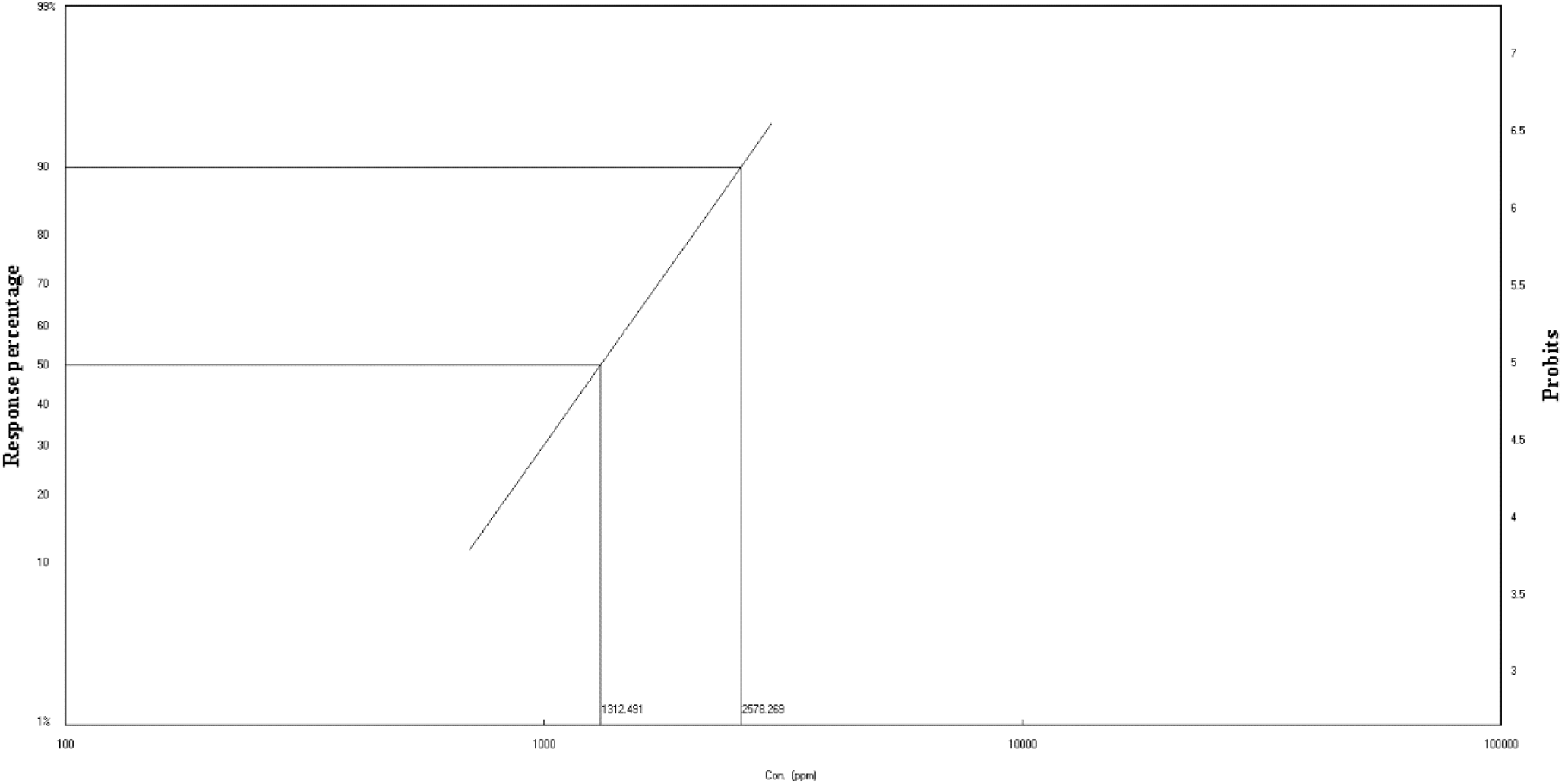
The LDP line of LC_50_ and LC_90_ against *A. aegypti*.

The results also showed malformation effects against treated larvae with boric acid compared to the control group (Fig. 2). These malformations manifested in Albino pupa (Fig. 3, b), Pigmentation (Fig. 3, c, h, i, n, o, q, v), Nick elongation (Fig. 3, v), Segments contraction (Fig. 3, h, i, p), Deformed cuticles (Fig. 3, j, k, r, x, z), Opaque swelling on the thorax (Fig. 3, f, g, n, o, r, t), Black coloration at the posterior end (Fig. 3, e, p, t, u), Larval-pupal intermediate (Fig. 3, d, l, m, s, u, w). The deformations activity of the boric acid can be attributed to its chemical components, which include a derivative of minerals used for insect control ^[5]^.

**Fig 2:**
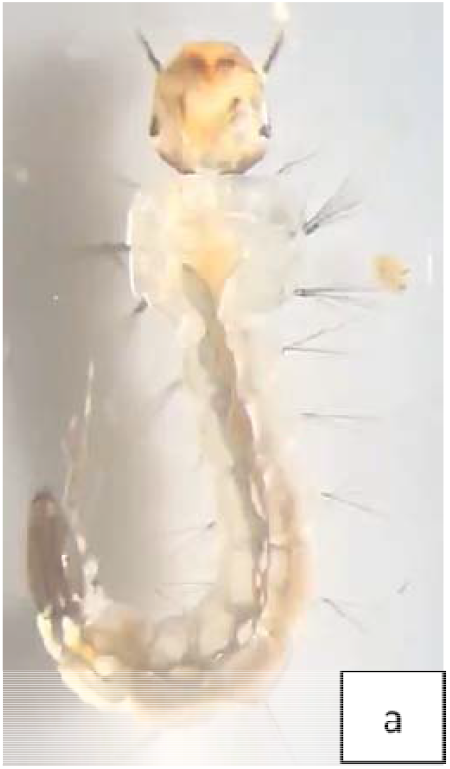
Normal 4^th^ larval stage of *A. aegypti*

**Fig 3:**
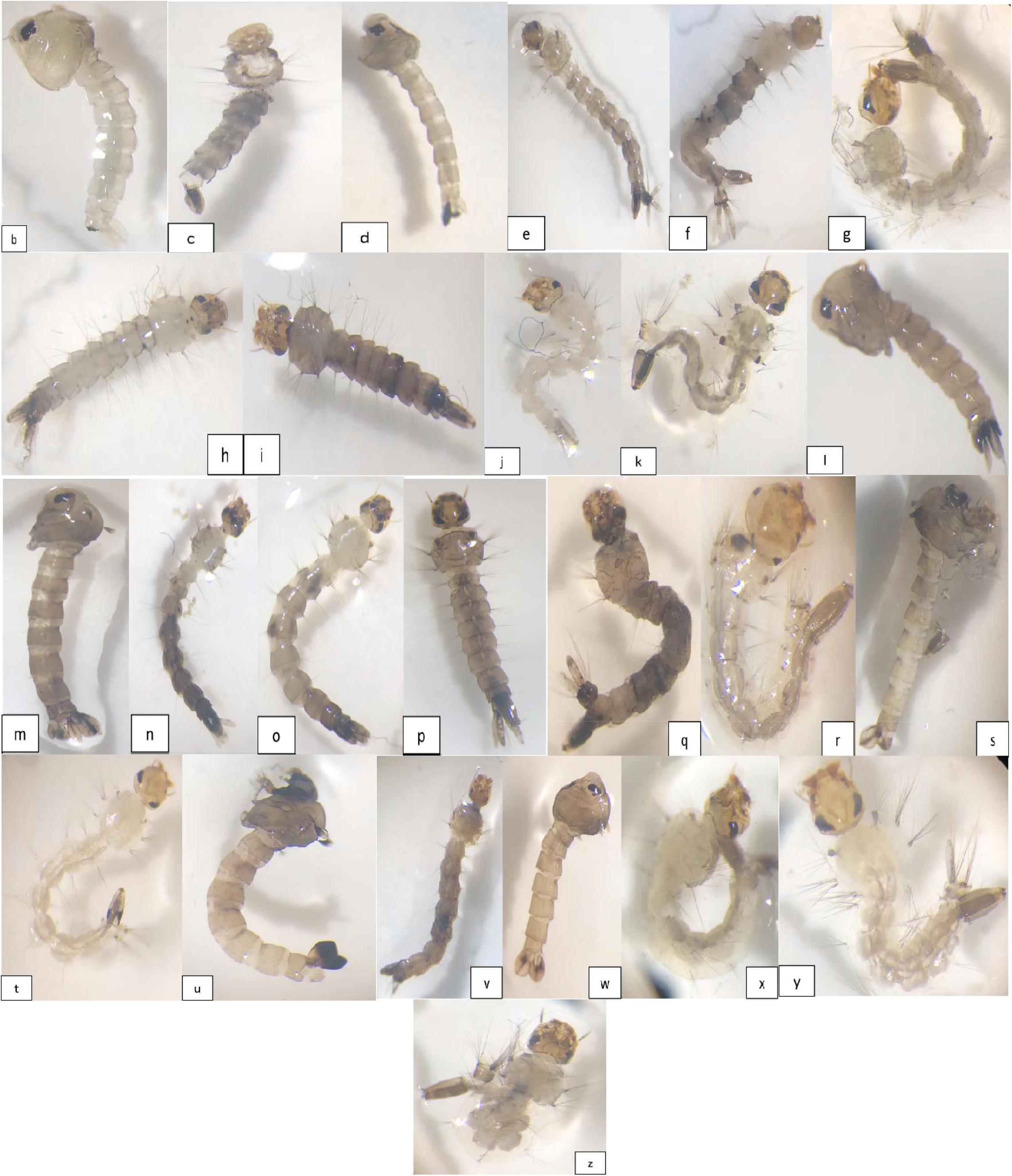
Some abnormalities of *A. aegypti* larval. b: Albino pupa, c: Pigmentation, d: Larval-pupal intermediate, e: Black coloration at the posterior end, f&g: Opaque swelling on the thorax, h&i: Pigmentation and segments contraction, j&k: Deformed cuticles, l&m: Larval-pupal intermediate, n&o: Pigmentation and opaque swelling on the thorax, p: Segments contraction and black coloration at the posterior end, q: Pigmentation, r: Deformed cuticles and opaque swelling on the thorax, s: Larval-pupal intermediate, t: Opaque swelling on the thorax and black coloration at the posterior end, u: Larval-pupal intermediate and black coloration at the posterior end, v: Pigmentation and nick elongation, w: Larval-pupal intermediate, x-z: Deformed cuticles and opaque swelling on the thorax.

## Conclusions

Boric acid can be effective in controlling the larval stages of *A. aegypti*. However, it showed mortality and malformation effects might be due to its chemical components including derivatives of minerals that can be used for insect control. Further, the use of boric acid as a safe method for controlling larval stages of *A. aegypti* can be recommended as it is available, cost-effective, and less harmful to humans, animals, and the environment.

## Author contribution

Dr. Somia Sharawi conceived and designed the study conducted research, provided research materials, and collected and organized data. Also, analyzed and interpreted data, wrote the initial and final draft of the article, and provided logistic support. The author has critically reviewed and approved the final draft and is responsible for the content and similarity index of the manuscript

## Ethical approval

There is no ethical issue

